# The C-terminus is critical for the degradation of substrates by the *Pseudomonas aeruginosa* CtpA protease

**DOI:** 10.1101/2020.03.31.019315

**Authors:** Sammi Chung, Andrew J. Darwin

## Abstract

Bacterial carboxyl-terminal processing proteases (CTPs) are widely conserved and have been linked to important processes including signal transduction, cell wall metabolism, and virulence. However, the features that target proteins for CTP-dependent cleavage are unclear. Studies of the *Escherichia coli* CTP Prc suggested that it cleaves proteins with non-polar and/or structurally unconstrained C-termini, but it is not clear if this applies broadly. *Pseudomonas aeruginosa* has a divergent CTP, CtpA, which is required for virulence. CtpA works in complex with the outer membrane lipoprotein LbcA to degrade cell wall hydrolases. Here, we investigated if the C-termini of two non-homologous CtpA substrates are important for their degradation. We determined that these substrates have extended C-termini, compared to their closest *E. coli* homologs. Removing seven amino acids from these extensions was sufficient to inhibit their degradation by CtpA both *in vivo* and *in vitro*. Degradation of one truncated substrate was restored by adding the C-terminus from the other, but not by adding an unrelated sequence. However, modification of the C-terminus of non-substrates, by adding the C-terminal amino acids from a substrate, did not cause their degradation by CtpA. Therefore, the C-termini of CtpA substrates are required but not sufficient for degradation. Although C-terminal truncated substrates were not degraded, they still associated with the LbcA•CtpA complex *in vivo*. Therefore, degradation of a protein by CtpA requires a C-terminal-independent interaction with the LbcA•CtpA complex, followed by C-terminal-dependent degradation, perhaps because CtpA must initiate cleavage at a specific C-terminal site.

**IMPORTANCE:** Carboxyl-terminal processing proteases (CTPs) are found in all three domains of life, but exactly how they work is poorly understood, including how they recognize substrates. Bacterial CTPs have been associated with virulence, including CtpA of *Pseudomonas aeruginosa*, which works in complex with the outer membrane lipoprotein LbcA to degrade potentially dangerous peptidoglycan hydrolases. We report an important advance by revealing that degradation by CtpA requires at least two separable phenomena, and that one of them depends on information encoded in the substrate C-terminus. A C-terminal-independent association with the LbcA•CtpA complex is followed by C-terminal-dependent cleavage by CtpA. Increased understanding of how CTPs target proteins is significant, due to their links to virulence, peptidoglycan remodeling, and other important processes.

## INTRODUCTION

*Pseudomonas aeruginosa* is a widespread Gram-negative bacterium and a frequent cause of serious opportunistic human infections (1). Like all bacterial pathogens, many *P. aeruginosa* virulence factors are assembled in its cell envelope, or must pass through the envelope on their way out of the bacterial cell. These include type II and III secretion systems, their exported substrates, pili, and the extracellular polysaccharide alginate, which plays an important role during chronic lung infections of cystic fibrosis patients (1-3). The successful production and function of these virulence factors is presumably impacted by the normal physiological functions that produce, maintain and remodel the major components of the cell envelope (4).

Proteolysis is an important process in the bacterial cell envelope. It ranges from discrete processing events during protein export, assembly, and signal transduction, to the complete degradation of proteins, which is especially important for misfolded or otherwise dangerous proteins (e.g. 5, 6, 7). One family of proteases found in the bacterial cell envelope is the carboxyl-terminal processing proteases (CTPs), which also occur in archaea and eukaryotes. These enzymes belong to the S41 family of serine proteases, and they can cleave a substrate once for processing, or degrade them completely (8-11). The CTP name is derived from early findings that these proteases often cleave close to the C-terminus of their substrate. However, at least one member of the CTP family in *Xanthomonas campestris* has now been shown to cleave close to the N-terminus of its substrate (12).

Recently, it has emerged that one role played by Gram-negative bacterial CTPs is to control potentially dangerous cell wall cross-link hydrolases by degrading them (13, 14). This is likely to be a widespread phenomenon, because it occurs in the divergent species *Escherichia coli* and *P. aeruginosa*. In *E. coli*, the Prc protease degrades the cell wall cross-link hydrolase MepS, and in *P. aeruginosa* CtpA degrades at least four predicted cell wall cross-link hydrolases (13, 14). Both of these CTPs form complexes with an outer membrane lipoprotein that is required for their proteolytic function, NlpI in *E. coli* and LbcA in *P. aeruginosa* (13, 14). However, despite their obvious similarities, Prc and CtpA are not orthologs. Prc is much larger than CtpA, and it is a member of the CTP-1 subfamily, whereas CtpA is in the divergent CTP-3 subfamily (15-17). Furthermore, although their lipoprotein binding partners NlpI and LbcA both contain the short, degenerate tetratricopeptide repeat motifs, they do not share obvious primary sequence similarity and are very different in size.

Exactly how CTPs recognize their substrates is unclear. In *E. coli*, Prc has been suggested to target proteins with non-polar and/or structurally unconstrained C-termini (10, 18-20). However, nothing is known about whether or not the C-terminus of a substrate is important during cleavage by the CtpA protease of *P. aeruginosa*. The significant differences between Prc and CtpA mean that what is true for Prc cannot be assumed to be true for CtpA. In particular, they might not share similar substrate recognition requirements. Indeed, the substrates of CtpA do not have non-polar C-termini (14). Knowing the features required for substrate cleavage is an important step towards understanding how CtpA-dependent proteolysis is controlled in *P. aeruginosa*. Therefore, we have investigated the role of the substrate C-terminus. Our results reveal that the extreme C-terminus is a critical determinant of degradation both *in vivo* and *in vitro*, and that the C-termini of two unrelated substrates are functionally interchangeable for degradation. However, the C-terminal motif is not sufficient for degradation, which suggests that it is only one of two or more checkpoints that determine if a protein will be degraded by CtpA.

## RESULTS

### Truncation of the C-terminus protects substrates from degradation by CtpA *in vivo*

The first CtpA substrate to be discovered was PA0667, which we named MepM due to its homology with the *E. coli* MepM peptidoglycan cross-link hydrolase (14, 21). *P. aeruginosa* MepM has a short C-terminal extension that is not present in *E. coli* MepM, but is conserved in other *Pseudomonas* species, all of which also have CtpA (Fig. 1A). We hypothesized that this extension might be important for CtpA-dependent degradation. To test this, we constructed *araBp-mepM* expression plasmids encoding MepM with C-terminal truncations and determined their steady state levels in *ctpA*^+^ and Δ*ctpA* strains. C-terminal truncation increased the amount of MepM in *ctpA*^+^ cells, suggesting protection from CtpA-dependent degradation (Fig. 1C). The Δ6 truncation (C-terminal six amino acids removed) reproducibly had a slightly higher level in Δ*ctpA* compared to *ctpA*^+^ strains, whereas longer truncations (Δ7-Δ9) consistently had a similar level in both strains (Fig. 1C and data not shown). The Δ8 truncation reduced the amount of MepM even in the Δ*ctpA* strain, perhaps due to CtpA-independent destabilization (Fig. 1C and data not shown). Therefore, from all these data we concluded that removing approximately seven C-terminal amino acids was sufficient to protect MepM from CtpA-dependent degradation *in vivo*, without otherwise destabilizing the protein.

**FIG 1.**
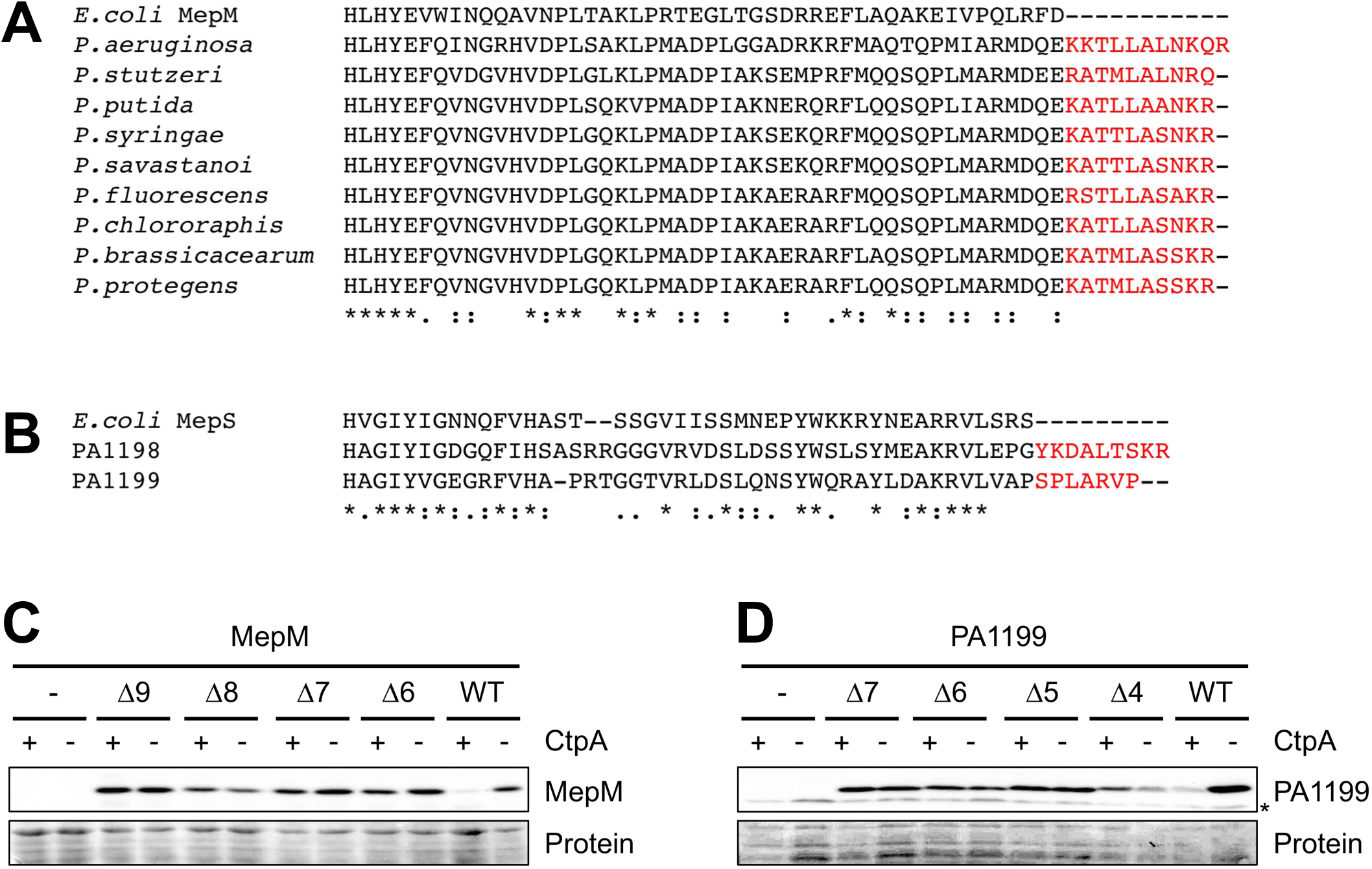
Truncation of the C-termini of MepM and PA1199 protects them from degradation by CtpA *in vivo*. (A) CLUSTAL Omega alignment of the C-termini of MepM proteins from *E. coli* and *Pseudomonas* species. (B) CLUSTAL Omega alignment of the C-termini of *E. coli* MepS, and PA1198 and PA1199 from *P. aeruginosa*. For panels A and B, amino acids in red represent C-terminal extensions in the *Pseudomonas* proteins. (C) MepM immunoblot analysis of equivalent amounts of whole-cell lysates of *ctpA*^+^ and Δ*ctpA* strains. (D) PA1199 immunoblot analysis of equivalent amounts of whole-cell lysates of *ctpA*^+^ and Δ*ctpA* strains (an asterisk indicates a protein cross-reactive with the PA1199 antiserum). For panels C and D, strains contained an arabinose-inducible expression plasmid encoding wild type MepM or PA1199 (WT), or derivates with the indicated number of amino acids removed from the C-terminus, and were grown in medium containing arabinose. MepM or PA1199 were detected with polyclonal antisera, and a Ponceau S total protein stain of the same region of the nitrocellulose membrane is shown to document loading levels.

MepM and another CtpA substrate, PA4404, are both members of the LytM/M23 peptidase family. However, the two other known substrates, PA1198 and PA1199, are in the NlpC/P60 peptidase family (14). PA1198 and PA1199 are over 50% identical to each other, and homologous to *E. coli* MepS. An alignment of *E. coli* MepS with PA1198 and PA1199 revealed that the *P. aeruginosa* proteins have extended C-termini (Fig. 1B). Therefore, we expanded our analysis to include one of these NlpC/P60 family substrates, PA1199. As with MepM, C-terminal truncations increased the amount of PA1199 in *ctpA*^+^ cells, suggesting protection from CtpA-dependent degradation (Fig. 1D). In this case, removing five C-terminal amino acids appeared sufficient to protect PA1199 from CtpA-dependent degradation *in vivo*, without otherwise destabilizing the protein. Together, all of these data suggest that the C-termini of MepM and PA1199 contain information that is needed for their degradation by CtpA *in vivo*.

### The C-termini of MepM and PA1199 are interchangeable for CtpA-dependent degradation

To extend our investigation we focused on the Δ7 truncations of MepM and PA1199, because both rendered the proteins equally abundant in *ctpA*^+^ and Δ*ctpA* strains (Fig. 1). The amino acids removed from each protein are not obviously similar (LALNKQR for MepM, and SPLARVP for PA1199, Fig. 1). Nevertheless, we tested our conclusion that they contain specific information required for CtpA-dependent degradation, by exchanging them.

MepM-Δ7 was similarly abundant in *ctpA*^+^ and Δ*ctpA* strains as seen previously (Figs. 1 and 2). However, when the C-terminal seven amino acids of PA1199 were added onto the C-terminus of MepM-Δ7 it behaved indistinguishably from wild type MepM, suggesting that its degradation by CtpA was restored (Fig. 2A). Similarly, addition of the C-terminal seven amino acids of MepM onto the end of PA1199-Δ7 made it behave similarly to wild type PA1199 (Fig. 2B). These experiments further support the conclusion that the C-termini of CtpA substrates are required for their degradation *in vivo*, and also show that they can be exchanged between two different substrates.

**FIG 2.**
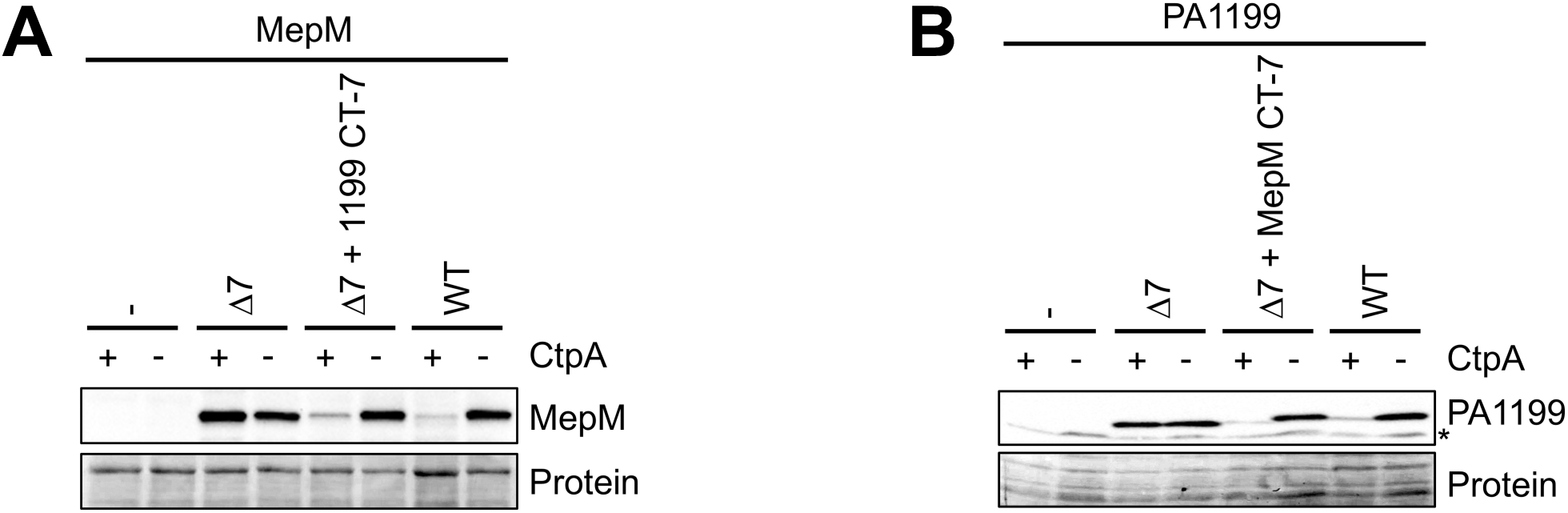
Substrate C-termini are interchangeable for degradation by CtpA. (A) MepM immunoblot analysis of equivalent amounts of whole-cell lysates of *ctpA*^+^ and Δ*ctpA* strains. Strains contained an arabinose-inducible expression plasmid encoding wild type MepM (WT), or derivates with seven amino acids removed from the C-terminus (Δ7), or with the C-terminal seven amino acids replaced by those from PA1199 (Δ7 + 1199 CT-7). (B) PA1199 immunoblot analysis of equivalent amounts of whole-cell lysates of *ctpA*^+^ and Δ*ctpA* strains (an asterisk indicates a protein cross-reactive with the PA1199 antiserum). Strains contained an arabinose-inducible expression plasmid encoding wild type PA1199 (WT), or derivates with seven amino acids removed from the C-terminus (Δ7), or with the C-terminal seven amino acids replaced by those from MepM (Δ7 + MepM CT-7). For both panels, strains were grown in medium containing arabinose. MepM or PA1199 were detected with polyclonal antisera, and a Ponceau S total protein stain of the same region of the nitrocellulose membrane is shown to document loading levels.

### Restoring the length of truncated substrates is not sufficient for CtpA-dependent degradation

CtpA-dependent degradation of two substrates truncated by seven amino acids was restored by adding back seven amino acids from a different substrate (Fig. 2). This raised the question of whether it is the length of the substrates that is critical, or if their C-termini contain sequence-dependent information. To distinguish between these possibilities, we generated a seven amino acid sequence randomly, AGEAGHL, and added it to the C-termini of the MepM-Δ7 and PA1199-Δ7 proteins. In contrast to the C-terminal swaps, adding these random seven amino acids to the truncated MepM and PA1199 proteins did not reduce their levels in a *ctpA*^+^ strain compared to a Δ*ctpA* strain (Fig, 3). Therefore, the C-terminal sequences of MepM and PA1199 contain sequence-specific information important for degradation by CtpA.

### The C-terminal motif of a CtpA substrate is not sufficient for CtpA-dependent degradation

In *E. coli*, it was reported that adding the C-terminal five amino acids of a Prc substrate onto the C-terminus of a non-substrate rendered it cleavable by Prc *in vivo* and *in vitro* (22). Our findings suggested that the C-terminal amino acids of CtpA substrates are also required for degradation, but they did not address if they are sufficient for degradation. Therefore, we tested this possibility next.

Besides the CtpA substrates MepM and PA4404, *P. aeruginosa* has a third member of the LytM/M23 peptidase family in its cell envelope that is predicted to be catalytically active, PA3787. However, PA3787 is not a CtpA substrate *in vivo* or *in vitro* (14). To test if the C-terminal amino acids of a substrate are sufficient for CtpA-dependent degradation, we added the C-terminal seven amino acids of MepM onto the C-terminus of PA3787. This hybrid protein was slightly less abundant than wild type PA3787 in both *ctpA*^+^ and Δ*ctpA* cells (Fig. 3A). However, it did not accumulate in a Δ*ctpA* strain compared to a *ctpA*^+^ strain. Therefore, the addition of the C-terminal seven amino acids from MepM did not make the related PA3787 a CtpA substrate *in vivo*.

**FIG 3.**
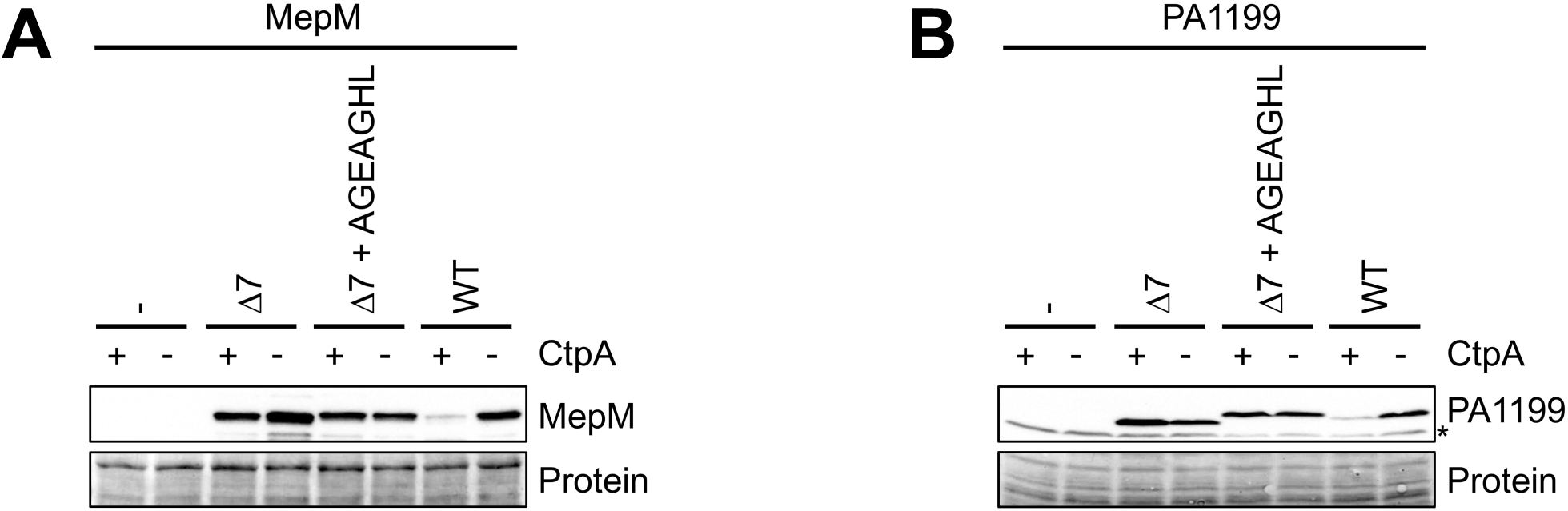
Restoring the length of truncated substrates is not sufficient for CtpA-dependent degradation. (A) MepM immunoblot analysis of equivalent amounts of whole-cell lysates of *ctpA*^+^ and Δ*ctpA* strains. Strains contained an arabinose-inducible expression plasmid encoding wild type MepM (WT), or derivates with seven amino acids removed from the C-terminus (Δ7), or with the C-terminal seven amino acids replaced by AGEAGHL. (B) PA1199 immunoblot analysis of equivalent amounts of whole-cell lysates of *ctpA*^+^ and Δ*ctpA* strains (an asterisk indicates a protein cross-reactive with the PA1199 antiserum). Strains contained an arabinose-inducible expression plasmid encoding wild type PA1199 (WT), or derivates with seven amino acids removed from the C-terminus (Δ7), or with the C-terminal seven amino acids replaced by AGEAGHL. For both panels, strains were grown in medium containing arabinose. MepM or PA1199 were detected with polyclonal antisera, and a Ponceau S total protein stain of the same region of the nitrocellulose membrane is shown to document loading levels.

PA3787 lacks the LysM peptidoglycan-binding domain that is found in MepM, making it a shorter protein with only 18% overall identity to MepM. Therefore, to test our conclusion more rigorously we extended our experiments to include *E. coli* MepM, which in comparison to PA3787, is a closer homolog of *P. aeruginosa* MepM. *P. aeruginosa and E. coli* MepM are similar in length, have the same domain organization, and are approximately 30% identical (not shown). Both have LysM peptidoglycan-binding and LytM/M23 peptidase domains, and their homology is distributed throughout their length (not shown). However, the C-terminal KKTKLALNKQR motif of *P. aeruginosa* MepM is absent from *E. coli* MepM (Fig. 1A). To further test if the C-terminal amino acids are sufficient for degradation by CtpA, we added the entire KKTKLALNKQR sequence onto the C-terminus of *E. coli* MepM. This was followed by a FLAG-tag sequence, because we did not have a reagent to detect *E. coli* MepM, and because a C-terminal FLAG tag does not affect the degradation of any CtpA substrate, including MepM (14). As expected, *P. aeruginosa* MepM-FLAG was undetectable in a *ctpA*^+^ strain, but abundant in a Δ*ctpA* mutant, consistent with its degradation by CtpA as in our previous report (Fig. 3B; ref. 14). In contrast, *E. coli* MepM-FLAG was equally abundant in both strains, suggesting that it is not degraded by CtpA. Furthermore, *E. coli* MepM-FLAG with the KKTKLALNKQR motif added was also equally abundant in both strains. Therefore, these experiments further support the conclusion that the C-terminus of a CtpA substrate is required, but not sufficient, for degradation by CtpA *in vivo*.

### Truncation of the C-terminus protects substrates from degradation by CtpA *in vitro*

It was possible that the C-terminal truncation mutations of MepM and PA1199 had an indirect effect *in vivo*, rather than rendering the proteins resistant to direct degradation by CtpA. One example would be if their localization was affected so that they became separated from CtpA. Therefore, we used an *in vitro* proteolysis assay to test our conclusion that the C-terminal amino acids are required for direct degradation by CtpA. The CtpA-binding partner LbcA, which increases CtpA activity, CtpA itself, and an inactive CtpA-S302A control (catalytic serine changed to alanine) were purified with C-terminal His_6_ tags, as before (14). However, MepM, MepM-Δ7, PA1199 and PA1199-Δ7 were purified with N-terminal His_6_ tags, so that their C-termini were unaltered, and identical to the proteins we had studied in the *in vivo* experiments.

When MepM was incubated with CtpA and LbcA, it was mostly degraded after one hour, and completely degraded after three hours (Fig. 5A). In contrast, MepM-Δ7 was resistant to degradation at both timepoints, with no degradation evident after 1 hour, and only slight degradation after three hours. PA1199 was degraded more quickly than MepM *in vitro*, such that it was degraded completely after 30 minutes (Fig. 5B). However, the PA1199-Δ7 protein was completely resistant to degradation after one hour (Fig. 5B), and we have also found that it remains resistant to degradation after three hours (data not shown). These experiments suggest that the C-terminal seven amino acids of MepM and PA1199 are required for their direct degradation by the LbcA**•**CtpA complex.

**FIG 4.**
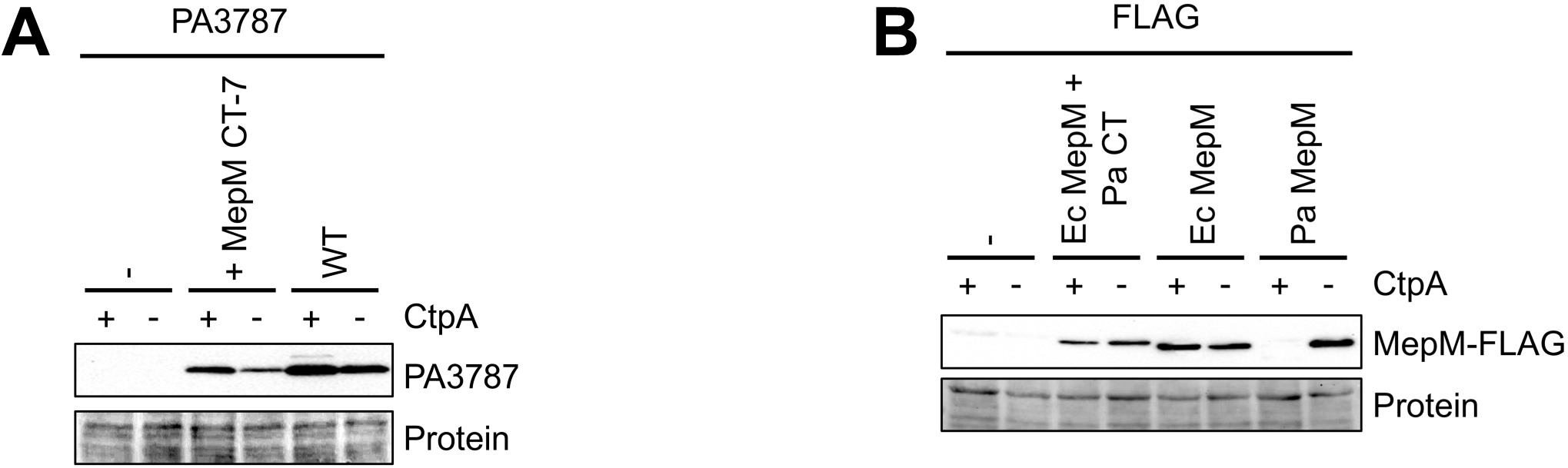
The C-terminal motif of a CtpA substrate is not sufficient for CtpA-dependent degradation. (A) PA3787 immunoblot analysis of equivalent amounts of whole-cell lysates of *ctpA*^+^ and Δ*ctpA* strains. Strains contained an arabinose-inducible expression plasmid encoding wild type PA3787 (WT), or a derivate with the seven C-terminal amino acids from MepM added to its C-terminus. (B) Anti-FLAG immunoblot analysis of equivalent amounts of whole-cell lysates of *ctpA*^+^ and Δ*ctpA* strains. Strains contained an arabinose-inducible expression plasmid encoding *P. aeruginosa* MepM (Pa), *E. coli* MepM (Ec), or *E. coli* MepM with the eleven C-terminal amino acids from P. aeruginosa MepM added to its C-terminus (+ Pa CT). The C-termini of all proteins terminated with the FLAG-tag sequence. For both panels, strains were grown in medium containing arabinose. Proteins were detected with ant-FLAG monoclonal antibodies, and a Ponceau S total protein stain of the same region of the nitrocellulose membrane is shown to document loading levels.

**FIG 5.**
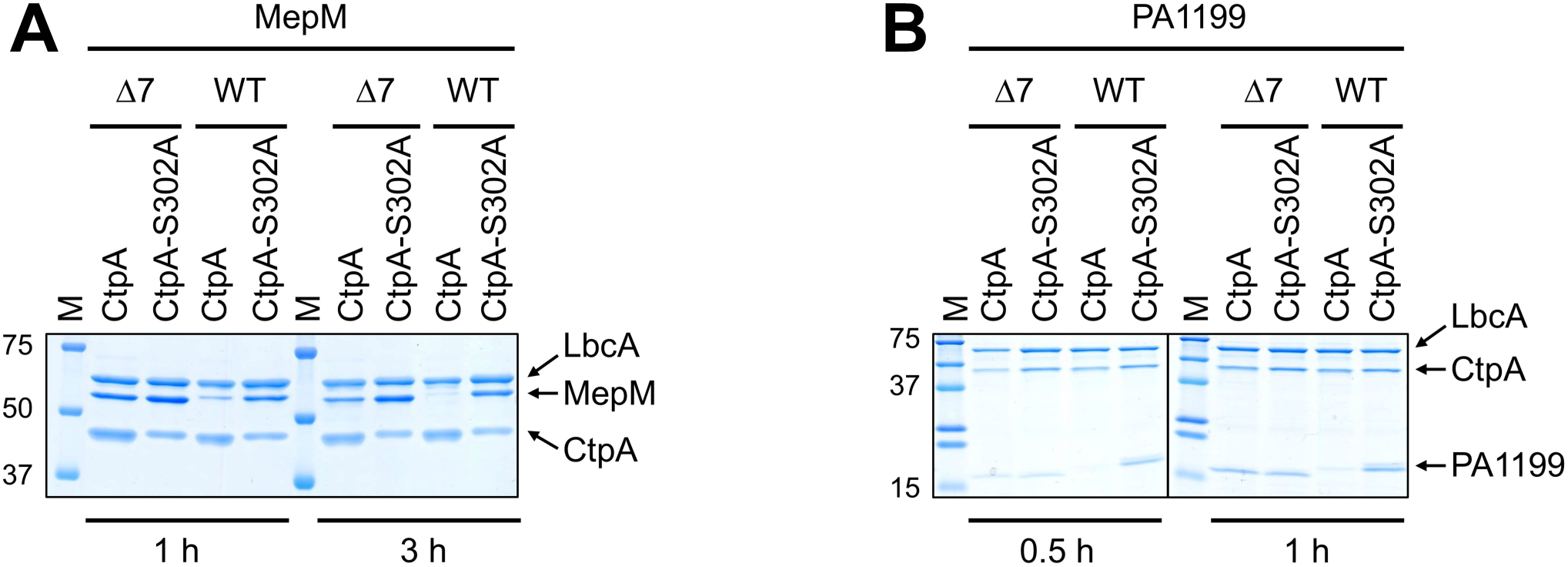
Truncation of the C-termini of MepM and PA1199 protects them from degradation by CtpA *in vitro*. (A) His_6_-MepM (WT) or His_6_-MepM-Δ7 proteins were incubated with LbcA-His_6_ and either active CtpA-His_6_ (CtpA) or inactive CtpA-S302A-His_6_ (CtpA-S302A), for 1 h or 3 h. Samples were separated by 10% SDS-PAGE and stained with ProtoBlue Safe (National Diagnostics). (B) His_6_-PA1199 (WT) or His_6_-PA1199-Δ7 proteins were incubated with LbcA-His_6_ and either active CtpA-His_6_ (CtpA) or inactive CtpA-S302A-His_6_ (CtpA-S302A), for 0.5 h or 1 h. Samples were analyzed on a single 16% SDS-PAGE gel, but the order of the left- and right-hand sides of the gel was reversed to construct the image, indicated by the vertical black line. For both panels, approximate kDa size of molecular-mass-marker proteins (M) are indicated on the left-hand side.

### Truncation of the C-terminus does not prevent substrates from associating with the LbcA•CtpA complex *in vivo*

MepM was discovered as a CtpA substrate because it copurified with the proteolytically inactive LbcA**•**CtpA-S302A complex after *in vivo* formaldehyde cross-linking (14). We considered the possibility that the truncated substrates might be resistant to degradation because they can no longer associate with the LbcA**•**CtpA complex. Alternatively, the truncated substrates might still associate with LbcA**•**CtpA, but their degradation is inhibited for another reason, perhaps because CtpA normally initiates cleavage from a missing C-terminal motif. To distinguish between these possibilities, we repeated the procedure that discovered MepM (14), to test if the MepM-Δ7 and PA1199-Δ7 proteins were still trapped by an inactive LbcA**•**CtpA-S302A complex.

Δ*ctpA* Δ*mepM* strains contained a plasmid encoding CtpA-S302A-His_6_, and a second plasmid encoding LbcA-FLAG and either MepM or MepM-Δ7. The LbcA**•**CtpA-S302A complex was purified using nickel agarose first to capture CtpA-S302A-His_6_, followed by anti-FLAG affinity gel to capture LbcA-FLAG. Coomassie staining of the samples after SDS-PAGE revealed that LbcA and CtpA-S302A purified abundantly, as expected (Fig. 6A). Confirmation that a LbcA•CtpA-S302A complex was isolated was provided by negative control purifications from strains that did not encode LbcA-FLAG, but still encoded CtpA-S302A-His_6_. Neither LbcA or CtpA-S302A purified abundantly from these controls (Fig. 6). Immunoblot analysis showed that both MepM and MepM-Δ7 were captured with the LbcA**•**CtpA-S302A complex, but not in the negative controls (Fig. 6A). A similar experiment to analyze PA1199 revealed that both PA1199 and PA1199-Δ7 were captured by the LbcA**•**CtpA-S302A complex as well (Fig. 6B). These results suggest that removal of the C-terminal seven amino acids from MepM or PA1199 does not significantly affect their association with the LbcA**•**CtpA complex *in vivo*. Therefore, it is likely that the C-termini of these substrates are required to initiate their degradation after they have been engaged by the proteolytic complex (see Discussion).

**FIG 6.**
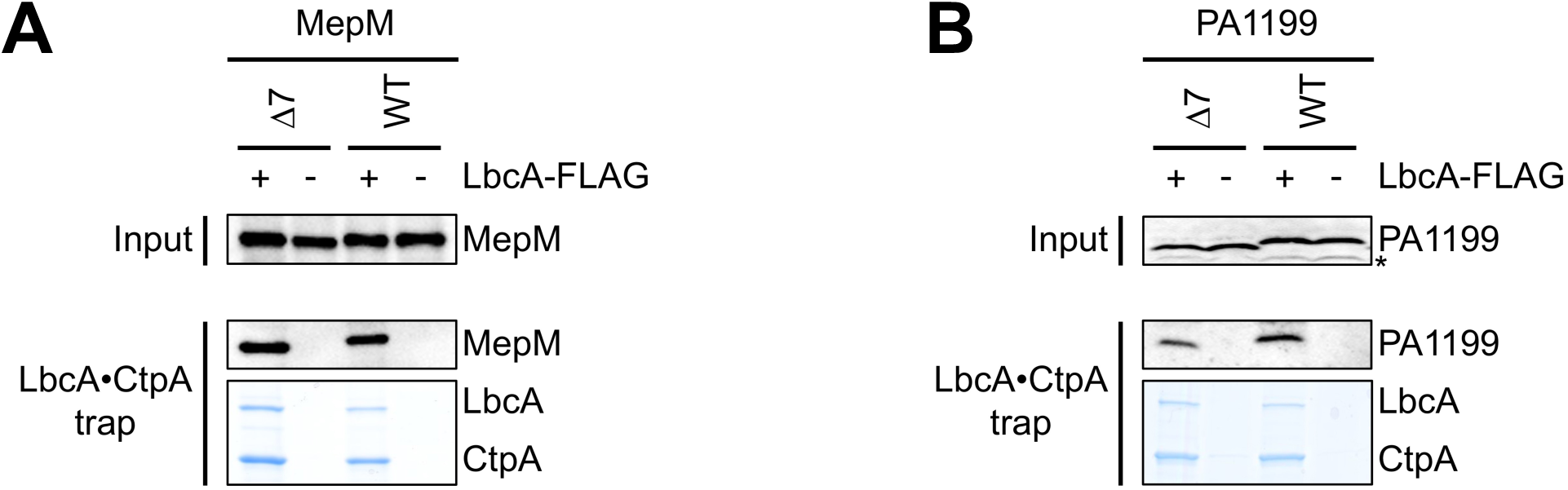
Truncation of the C-terminus does not prevent substrates from associating with the LbcA**•**CtpA complex *in vivo*. (A) Proteins were purified from detergent solubilized lysates of Δ*mepM* Δ*ctpA* strains, which contained one plasmid encoding CtpA-S302A-His_6_ and a second plasmid encoding MepM or MepM-Δ7 and LbcA-FLAG (+), or MepM or MepM-Δ7 only (-). (B) Proteins were purified from detergent solubilized lysates of Δ(PA1198-PA1199) Δ*ctpA* strains, which contained one plasmid encoding CtpA-S302A-His_6_ and a second plasmid encoding PA1199 or PA1199-Δ7 and LbcA-FLAG (+), or PA1199 or PA1199-Δ7 only (-). For both panels, tandem affinity purification of the LbcA**•**CtpA-S302A complex was done with nickel agarose followed by anti-FLAG M2 agarose resin. Input lysates (Input) and purified samples (LbcA**•**CtpA trap) were separated by SDS-PAGE and analyzed by anti-MepM (A) or anti-PA1199 (B) immunoblot (an asterisk indicates a protein cross-reactive with the PA1199 antiserum). Purified samples were also separated by SDS-PAGE and stained with ProtoBlue Safe (National Diagnostics) to monitor recovery of the LbcA**•**CtpA complex.

### A conserved amino acid at the -5 position is not essential for CtpA-dependent degradation

The sequence, charge, or hydrophobicity of the C-terminal amino acids of MepM and PA1199 do not have obvious similarity. However, an alignment of the C-terminal seven amino acids of all four CtpA substrates revealed that three of them have leucine as the fifth amino acid from their C-terminus (referred to as the -5 position) and the other one, PA4404, has alanine (Fig. 7A). These four CtpA substrates are LytM/M23 or NlpC/P60 peptidase family members. Two other predicted peptidoglycan cross-link hydrolase members of these families are not CtpA substrates, PA3787 and PA3472 (14). Neither of these has leucine or alanine at the -5 position (Fig. 7A). In ongoing work in our laboratory, we have identified a fifth CtpA substrate preliminarily, which also has leucine at the -5 position (D. Chakraborty, A. G. Sommerfield and A. J. Darwin, unpublished data). This means that three or four CtpA substrates have leucine at position -5 and one has alanine, whereas neither of two related non-substrates share this property. Therefore, we tested if the leucine at position -5 of MepM or PA1199 was important for their degradation by CtpA.

**FIG 7.**
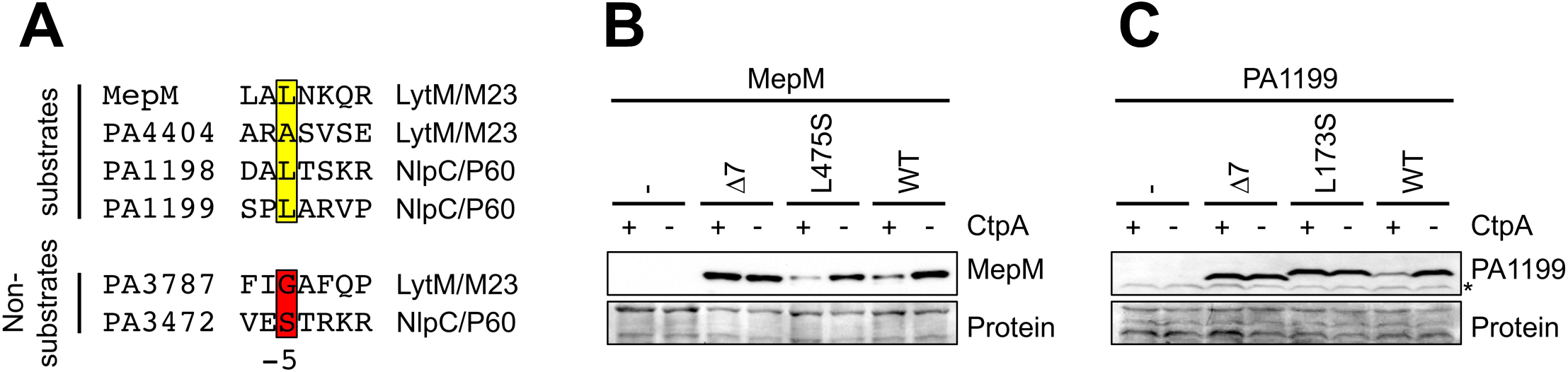
Mutation of the -5 position affects the CtpA-dependent stability of MepM and PA1199 differently. (A) Alignment of the C-terminal seven amino acids of predicted peptidoglycan cross-link hydrolases in the LytM/M23 or NlpC/P60 peptidase families that either are or are not CtpA substrates. The -5 positions of the substrates and non-substrates are highlighted yellow or red, respectively. (B) MepM immunoblot analysis of equivalent amounts of whole-cell lysates of *ctpA*^+^ and Δ*ctpA* strains. (C) PA1199 immunoblot analysis of equivalent amounts of whole-cell lysates of *ctpA*^+^ and Δ*ctpA* strains (an asterisk indicates a protein cross-reactive with the PA1199 antiserum). For panels B and C, strains were grown in medium containing arabinose and contained an arabinose-inducible expression plasmid encoding wild type MepM or PA1199 (WT), or derivates with the indicated amino acid substitution at the -5 position, or with seven amino acids removed from the C-terminus (Δ7). MepM or PA1199 were detected with polyclonal antisera, and a Ponceau S total protein stain of the same region of the nitrocellulose membrane is shown to document loading levels.

The non-substrate PA3472 has serine at the -5 position (Fig. 7A). Therefore, we made the conservative alanine to serine substitution at the -5 positions of MepM and PA1199. These substitutions had different effects on the steady state levels of the proteins. The MepM L475S mutant behaved indistinguishably from wild type MepM, showing that the leucine at position -5 is not required for its degradation by CtpA (Fig. 7B). In contrast, the PA1199 L173S mutant behaved indistinguishably from the PA1199-Δ7 truncation mutant, showing that its leucine at the -5 position is important for CtpA-dependent degradation. Taken together, these results suggest that while individual amino acids can influence CtpA-dependent destabilization *in vivo*, a conserved position-specific sequence signature for degradation is unlikely (see Discussion).

## DISCUSSION

CtpA is essential for *P. aeruginosa* T3SS function, for virulence in a mouse model of acute pneumonia, and it affects surface attachment (14, 17). These phenotypes are probably linked to the cell wall, because CtpA degrades peptidoglycan cross-link hydrolases (14). This means that CtpA is critical for fundamental cell envelope physiology, and for the ability of *P. aeruginosa* to cause disease. Therefore, it is important to understand all aspects of CtpA function, including the features of a protein that make it susceptible to CtpA-dependent proteolysis. The carboxyl-terminal processing protease (CTP) family was named because the first members studied were found to cleave close to the C-terminus of their substrates, either as a processing event, or to initiate degradation (e.g. 9, 10, 23). This suggests that substrate C-termini might contain information that is recognized by a CTP. Indeed, some CTPs, including *E. coli* Prc, have been proposed to target proteins with non-polar and/or structurally unconstrained C-termini (10, 18-20, 22, 24). However, the role of the substrate C-terminus has not been studied for most CTPs, and the same rules are unlikely to apply to all of them. In fact, *Xanthomonas campestris* Prc cleaves close to the N-terminus of a transmembrane substrate, and cannot rely on recognition features in the C-terminus, which is physically separated from Prc by the cytoplasmic membrane (12).

We have investigated if the C-termini of CtpA substrates play a role in their degradation, using one substrate in the LytM/M23 peptidase family (MepM), and one in the NlpC/P60 family (PA1199), as model substrates. Despite the fact that MepM and PA1199 are not homologous, in both cases their C-terminal amino acids were essential for degradation (Figs. 1 and 5). The C-terminal seven amino acids of MepM and PA1199 could also be exchanged without affecting their CtpA-dependent degradation *in vivo*, whereas substitution with an unrelated seven amino acids rendered the proteins resistant to degradation (Figs. 2-3). This suggests that the C-termini of CtpA substrates contain specific information required for degradation. However, the sequences of the seven C-terminal amino acids of MepM and PA1199 are not similar, and a comparison of all four CtpA substrates also failed to reveal obvious common C-terminal features (Fig. 7A). Notably, they are not predominantly non-polar, which is a feature that emerged from early research as something common amongst *E. coli* Prc substrates (20). We did notice that CtpA substrates have leucine or alanine at the -5 position, whereas two non-substrates in the LytM/M23 and NlpC/P60 peptidase families do not (Fig. 7A). However, although changing this leucine to serine protected PA1199 from CtpA-dependent degradation, it did not protect MepM (Fig. 7). From all of these observations, we conclude that the information required for CtpA-dependent degradation is not a conserved C-terminal amino acid sequence motif, but perhaps another property such as a structural feature.

An early study of *E. coli* Prc concluded that the C-terminal five amino acids of a substrate were sufficient for cleavage, because when added onto the C-terminus of a non-substrate it was cleaved by Prc (22). However, the addition of the *P. aeruginosa* MepM C-terminus onto two other LytM/M23 peptidases did not make either of them a CtpA substrate (Fig. 4). Therefore, the C-terminus of CtpA substrates is not sufficient for degradation. Later work on *E. coli* Prc revealed that it forms a complex with the outer membrane lipoprotein NlpI, which promotes Prc-dependent degradation of MepS *in vivo* and in *vitro* (13). Protein interaction and structure-function analysis suggested that NlpI binds to Prc and MepS independently, acting as a scaffold to bring protease and substrate together (10, 13). CtpA also has an outer membrane lipoprotein binding partner, LbcA, which promotes the degradation of all four CtpA substrates *in vivo* and *in vitro* (14). LbcA and NlpI are not homologous, but both contain tetratricopeptide repeats (TPR) that mediate the formation of multiprotein complexes (25). We have evidence that LbcA also acts as a scaffold, binding CtpA and its substrates independently (D. Chakraborty and A. J. Darwin, unpublished data). The MepM and PA1199 C-terminal truncation mutants still associated with the LbcA•CtpA complex *in vivo* (Fig. 6). This suggests that substrate C-termini are not involved in the association with LbcA. Therefore, the C-terminus of MepM is probably insufficient for degradation, because when transplanted onto a non-substrate it cannot provide a required association with the LbcA**•**CtpA complex. Interestingly, recent work indicated that cleavage of one proposed substrate of *E. coli* Prc, FtsI, is not helped by the NlpI binding partner of Prc *in vitro* (26). If something similar occurs *in vivo*, it would mean that NlpI promotes the cleavage of some substrates (MepS), but is not required for the cleavage of others (FtsI). That might explain why the transplantation of a Prc substrate C-terminus onto at least one non-substrate was sufficient for its degradation (22).

*P. aeruginosa* has a second CTP that is a close homolog of *E. coli* Prc, and has been named both Prc and AlgO. Early in this study we made an interesting observation about *P. aeruginosa* Prc that we are pursuing separately. When we began to analyze PA1199, some of truncation mutants had significantly reduced abundance in a Δ*ctpA* strain compared to the wild type protein (S. Chung and A. J. Darwin, unpublished data). *E. coli* Prc has been implicated in degrading proteins with aberrant C-termini (20, 27). Therefore, we reasoned that *P. aeruginosa* Prc might cleave some truncated PA1199 proteins due to their altered C-termini. In support of this, the abundance of the truncated PA1199 proteins was indistinguishable from full length PA1199 in Δ*ctpA* Δ*prc* strains (S. Chung and A. J. Darwin, unpublished data). For this reason, all of the *in vivo* analysis of PA1199 and its derivatives was done by comparing their abundance in Δ*prc ctpA*^+^ and Δ*prc* Δ*ctpA* strains, to eliminate any interference from Prc. Nevertheless, this suggests that *P. aeruginosa* Prc might play a role in protein quality control, which is consistent with the suggestion that Prc cleaves C-terminal truncated forms of MucA that arise in cystic fibrosis patients (28, 29). However, Prc does not degrade all proteins with aberrant C-termini because we did not detect any influence of Prc on truncated MepM proteins (S. Chung and A. J. Darwin, unpublished data).

Structural analysis of the *E. coli* NlpI•Prc complex, and a docking model of the C-terminal 12 amino acid peptide of its MepS substrate, suggests that the substrate C-terminus is bound by the PDZ domain of Prc (10). The Prc PDZ domain was proposed to recognize substrate C-termini with low specificity, because Prc degrades MepS with or without a C-terminal His_6_ tag, and will also degrade lysozyme *in vitro* if a disulfide bond in its C-terminus is broken by reduction (10). In contrast, our analysis suggests that there is specific recognition of the C-termini of CtpA substrates. However, all four CtpA substrates are still degraded *in vivo* when a FLAG tag is added onto their C-terminus, and *in vitro* with C-terminal His_6_ tags (14). This can still be reconciled with our conclusion that substrate C-termini are recognized specifically. We hypothesize that the native C-terminal amino acids of CtpA substrates can still make a required specific interaction with CtpA even if non-native amino acids have been added onto them.

In summary, this work has shown that the C-termini of *P. aeruginosa* CtpA substrates contain specific information required, but not sufficient, for degradation. We hypothesize that the cleavage of a protein by CtpA requires at least two phenomena: (1) association of the substrate with the LbcA•CtpA complex (most likely with LbcA) and (2) a specific recognition of the substrate C-terminus by CtpA. However, any interaction with LbcA might have to occur in a specific way, perhaps with one or more specific TPR motif(s), of the eleven that are present in LbcA. This is because we have evidence that LbcA might associate with some proteins that are not cleaved by CtpA, including PA3787 (ref. 14 and D. Chakraborty and A. J. Darwin, unpublished data). The NlpI partner of *E. coli* Prc has also been proposed to interact with some non-Prc substrates (30). Perhaps CtpA substrates and non-substrates interact differently with LbcA, possibly engaging different TPR motifs. Regardless, specific engagement of a substrate by LbcA, and then recognition of its C-terminus by CtpA, would be followed by the first cleavage event. Degradation might then proceed in a non-specific manner that does not require specific recognition of the new C-terminus generated after this first cleavage event. This might occur similarly to the lever-like mechanism proposed for *E. coli* Prc, which feeds the substrate into the Prc active site in a C- to N-terminal direction after each cleavage event (10). However, CtpA and Prc are in different CTP subfamilies and their lipoprotein partners are not homologous. Therefore, the details of the organization and function of the LbcA**•**CtpA complex are likely to diverge from those of *E. coli* NlpI**•**CtpA. The goal of future work will be to uncover exactly what those details are.

## MATERIALS AND METHODS

### Bacterial strains, plasmids and routine growth

Strains and plasmids are listed in Table 1. Bacteria were grown routinely in Luria-Bertani (LB) broth, composed of 1% (w/v) Tryptone, 0.5% (w/v) yeast extract and 1% (w/v) NaCl, or on LB agar, at 30°C or 37°C. *P. aeruginosa* was occasionally grown on Vogel-Bonner minimal agar (31).

**TABLE 1.**
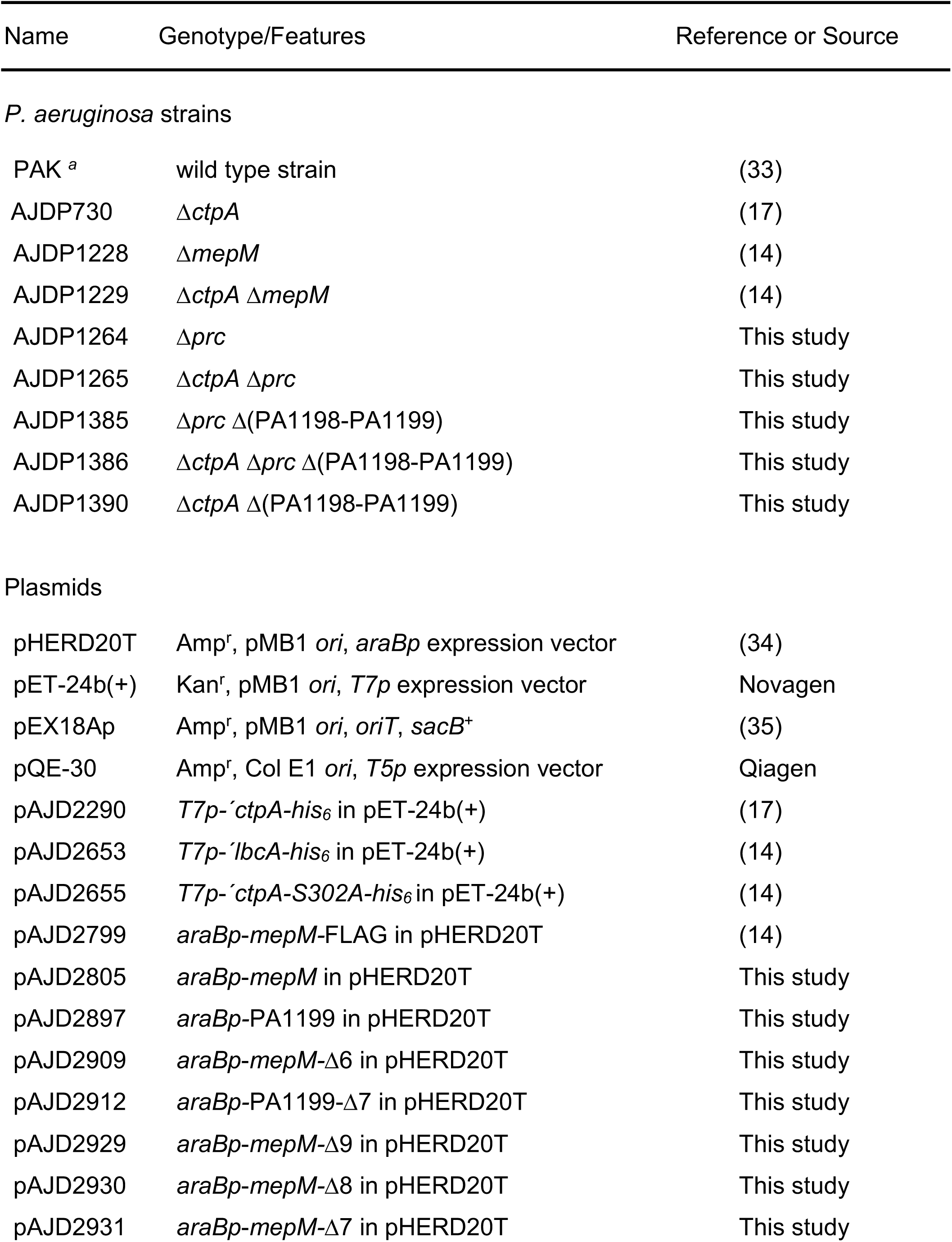
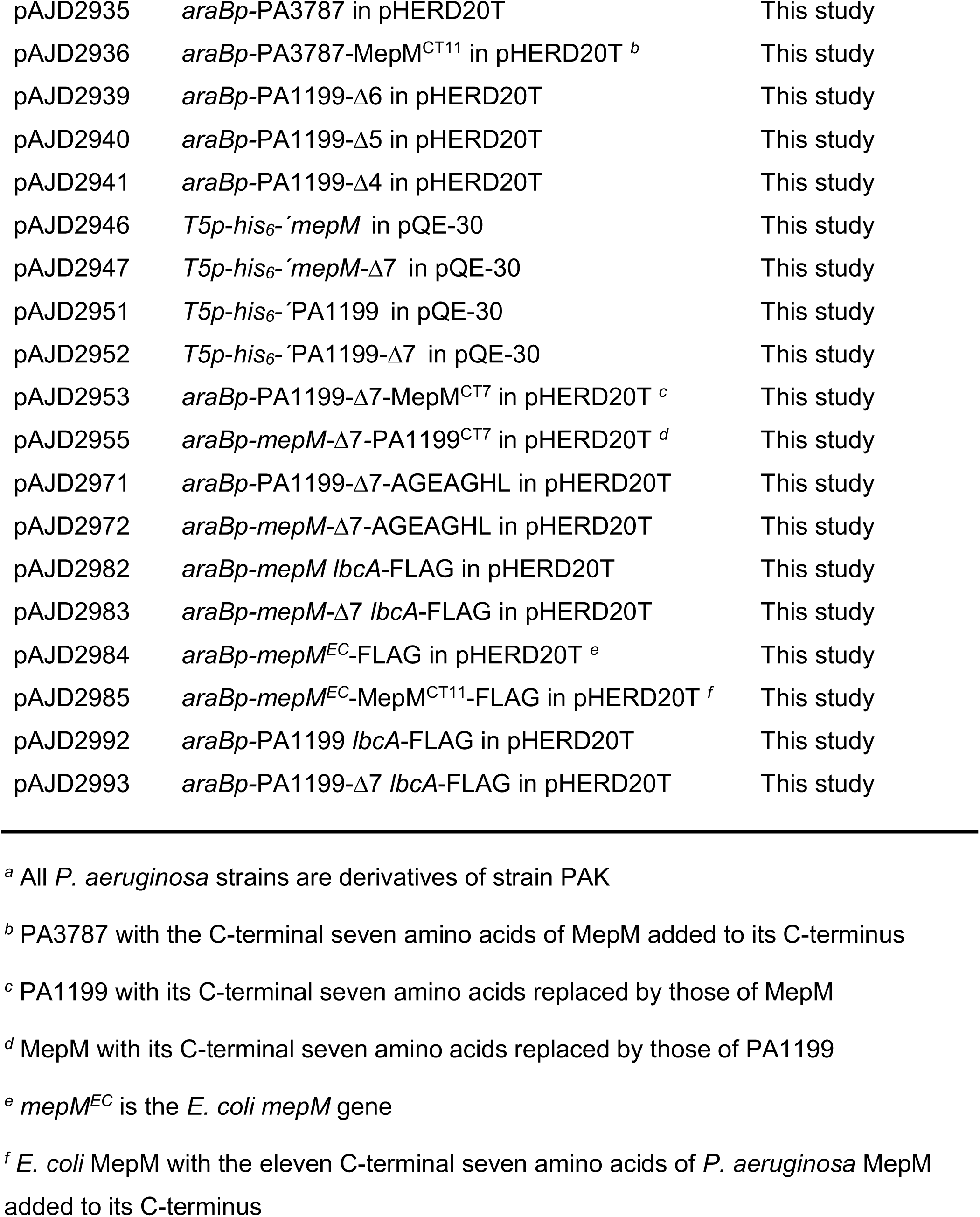
Strains and plasmids.

### Strain constructions

To construct Δ*prc* and Δ(PA1198-PA1199) in frame deletion mutants, two fragments of ∼ 0.5 kb each corresponding to the regions flanking the deletion site were amplified by PCR and cloned into pEX18Ap. The plasmids were integrated into the *P. aeruginosa* chromosome after conjugation from *E. coli* strain SM10 (32) and sucrose resistant, carbenicillin sensitive segregants were isolated on LB agar containing 10% sucrose. Deletions were verified by genomic PCR analysis.

### Plasmid constructions

*araBp* expression plasmids encoding MepM, PA1199 or PA3787 were constructed by amplifying the genes from *P. aeruginosa* chromosomal DNA using one primer that annealed ∼ 30 bp upstream of the start codon and a second primer that annealed immediately downstream of the stop codon. Plasmids encoding C-terminal truncated derivatives were constructed similarly, except that the downstream primers annealed within the gene and incorporated a premature stop codon. Plasmids encoding MepM or PA1199 with their C-terminal seven amino acids exchanged, or replaced by AGEAGHL, were constructed using downstream primers that annealed 21 bp upstream of the stop codon and incorporated a region encoding the final seven amino acids of MepM, PA1199, or the AGEAGHL sequence, followed by a stop codon. To construct a plasmid encoding PA3787 with the C-terminal seven amino acids of MepM added, the downstream primer annealed immediately upstream of the PA3787 stop codon and incorporated a region encoding the C-terminus of MepM followed by a stop codon. Plasmids encoding *E. coli* MepM-FLAG were constructed by amplifying *mepM* from *E. coli* strain MG1655 chromosomal DNA. The forward primer annealed immediately upstream of the start codon and incorporated the ribosome binding site from pQE-30, and the reverse primer annealed immediately upstream of the stop codon and incorporated a region encoding the FLAG tag only, or the C-terminal eleven amino acids of *P. aeruginosa* MepM followed by a FLAG tag, and a stop codon. In all cases, the amplified fragments were cloned into pHERD20T using restriction sites added to the fragments by the amplification primers.

For the LbcA**•**CtpA-S302A trap experiments, *araBp* expression plasmids were constructed to encode MepM or PA1199 full length or C-terminal truncated proteins, as well as LbcA-FLAG. *lbcA* was amplified from *P. aeruginosa* chromosomal DNA using a primer that annealed ∼ 40 bp upstream of the start codon and a primer that annealed immediately upstream of the stop codon and incorporated a region encoding the FLAG tag followed by a stop codon. This was cloned as an XbaI-HindIII fragment (restriction sites incorporated by the amplification primers) immediately downstream of the *mepM* or PA1199 genes in the expression plasmids described above.

pET-24b(+) derivatives used for overproduction and purification of LbcA-His_6_, CtpA-His_6_ and CtpA-S302A-His_6_ were described previously (14, 17). For overproduction and purification of His_6_-MepM or His_6_-PA1199 full length and C-terminal truncated proteins, the genes were amplified without their predicted N-terminal signal sequences and cloned into pQE-30 as BamHI-HindIII fragments.

### Determination of protein abundance *in vivo*

Saturated cultures were diluted into 5 ml of LB broth, containing 150 µg/ml carbenicillin and 0.02% (w/v) arabinose, in 18 mm diameter test tubes so that the initial OD 600 nm was 0.05. The cultures were grown on a roller drum at 37°C for 5 h. Cells were collected by centrifugation and resuspended in SDS-PAGE sample buffer at equal concentrations (based on the culture OD 600 nm) before being analyzed by immunoblot.

### Polyclonal antisera and immunoblotting

Proteins were separated by SDS-PAGE and transferred to a nitrocellulose membrane by semi-dry electroblotting. For analysis of total cell lysates, approximately equal loading and transfer was confirmed by total protein staining of the nitrocellulose membrane with Ponceau S (Amresco). Chemiluminescent detection followed incubation with polyclonal antiserum or monoclonal antibody, then goat anti-rabbit IgG (Sigma) or goat anti-mouse IgG (Sigma) horseradish peroxidase conjugates used at the manufacturers recommended dilution. The primary anti-FLAG M2 (Sigma) antibody was diluted 5,000-fold, and all polyclonal antisera used here have been described previously (14).

### Protein purification and *in vitro* proteolysis assay

LbcA-His_6_, CtpA-His_6_ and CtpA-S302A-His_6_ were encoded by pET-24b(+) derivatives in *E. coli* ER2566 (NEB). His_6_-MepM and His_6_-PA1199 full length and C-terminal truncated proteins were encoded by pQE-30 derivatives in *E. coli* M15 containing plasmid pREP4 to produce the LacI repressor (Qiagen). These strains were grown in 1L LB broth at 37°C with aeration until the OD 600 nm was 0.6-1.0. Protein production was induced by adding 1 mM IPTG and incubating for 3-4 h at 37°C (LbcA-His_6_, CtpA-His_6_ and CtpA-S302A-His_6_) or at 30°C (His_6_-MepM and His_6_-PA1199), with aeration. Proteins were purified under native conditions by NTA agarose affinity chromatography in buffer containing 50 mM NaH_2_PO_4_ and 300 mM NaCl, as recommended by the manufacturer (Qiagen). LbcA-His_6_, His_6_-MepM and His_6_-PA1199 proteins were eluted in fractions using 50 mM NaH_2_PO_4_ and 300 mM NaCl buffer containing increasing concentrations of imidazole (50 - 250 mM). Samples of each fraction were separated by SDS-PAGE, stained with ProtoBlue Safe (National Diagnostics), and 2-3 fractions judged to have the highest purity were combined, supplemented with 10% (w/v) glycerol and stored at -70°C. The same fractions were used for His_6_-MepM and for His_6_-MepM-Δ7, and the same fractions were used for His_6_-PA1199 and for His_6_-PA1199-Δ7. CtpA-His_6_ and CtpA-S302A-His_6_ were eluted similarly, but after combining fractions the proteins were concentrated ∼ 10-fold using Amicon Ultra-4 centrifuge filter devices (10 kDa cutoff), and then supplemented with 50% Protein Stabilizing Cocktail (ThermoFisher Scientific) before storing at -70°C. All *In vitro* proteolysis reactions contained approximately 2 µM of LbcA and CtpA or CtpA-S302A. MepM proteins were also used at 2 µM, but the smaller PA1199 proteins were used at approximately 15 µM to facilitate visualization after staining. Reactions were incubated at 37°C for 0.5 – 3 h, terminated by adding SDS-PAGE sample buffer and boiling, separated by SDS-PAGE, and stained with ProtoBlue Safe (National Diagnostics).

### Tandem affinity purification LbcA-FLAG•CtpA-S302A-His_6_ complex

Strains were grown to saturation, diluted to an OD 600nm of 0.05 in 400 ml of LB broth containing 5 mM EGTA, and shaken at 200 rpm for 2.5 h at 37°C. The cultures were supplemented with 0.02% (w/v) arabinose and 1 mM IPTG and shaken at 200 rpm for a further 3 h at 37°C. Cells from the equivalent of 200 ml of culture at OD 600nm of 1 were collected by centrifugation. The pellet was washed with cold 10 mM potassium phosphate buffer pH 8.0 and resuspended to an OD 600 nm of 5 in the same buffer. 1% formaldehyde was added followed by incubation at room temperature for 30 min. 0.3 M Tris-HCl pH 7.5 was added to quench and the cells were collected by centrifugation. Pellets were resuspended in 3 ml lysis buffer (50 mM Tris-HCl pH 7.5, 150 mM NaCl, 5 mM Imidazole), and Roche complete protease inhibitors (2x concentration), 1 µg/ml DNaseI, 1 µg/ml RNase, and 1 mg/ml lysozyme were added. Cells were disrupted by sonication, then 1% *n*-dodecyl β-D-maltoside (DDM) was added followed by incubation with rotation for 30 min at 4°C. Insoluble material was removed by centrifugation at 13,000 x *g* for 30 min at 4°C. 500 µl of nickel-NTA agarose in lysis buffer was added to the supernatant, followed by incubation with rotation for 50 min at 4°C. The resin was collected in a drip column and washed with 8 ml lysis buffer, then 8 ml lysis buffer containing 20 mM imidazole. Proteins were eluted in 1 ml lysis buffer containing 250 mM imidazole, mixed with 40 µl anti-FLAG M2 agarose resin (Sigma) in TBS (10 mM Tris-HCl pH 7.5, 150 mM NaCl), and incubated with rotation for 2 h at 4°C. A 1 ml spin column (Pierce 69725) was used to wash the resin seven times with 500 µl TBS. Proteins were eluted by adding 100 µl of 200 µg/ml 3xFLAG peptide (Sigma) in TBS and incubating with rotation at 4°C for 30 min.

## ACKNOWLEDGEMENTS

This study was supported by award number R01AI136901 from the National Institute of Allergy and Infectious Diseases (NIAID). The content is solely the responsibility of the authors and does not necessarily represent the official views of the NIAID or the National Institutes of Health.

We thank Dolonchapa Chakraborty for providing advice about the protein purification and *in vitro* proteolysis assays.

